# THE MODEL OF *PPARγ* DOWNREGULATED SIGNALING IN PSORIASIS

**DOI:** 10.1101/2020.09.01.274753

**Authors:** Vladimir Sobolev, Anastasia Nesterova, Anna Soboleva, Evgenia Dvoriankova, Anastas Piruzyan, Dzerassa Mildzikhova, Irina Korsunskaya, Oxana Svitich

**Affiliations:** I. Mechnikov Research Institute for Vaccines and Sera RAMS, Russian Federation, 105064, Moscow, Malyy Kazennyy per., 5; Centre of Theoretical Problems of Physico-Chemical Pharmacology, Russian Academy of Sciences, Russian Academy of Sciences, Russian Federation, 119334, Moscow, Kosygin str., 4; Life Science Research and Development Department, Elsevier Inc., Rockville, USA

## Abstract

Interactions of genes in intersecting signaling pathways, as well as environmental influences, are required for the development of psoriasis. Peroxisome proliferator-activated receptor gamma (*PPARγ*) is a nuclear receptor and transcription factor which inhibits the expression of many proinflammatory genes. We tested the hypothesis that low levels of *PPARγ* expression promote the development of psoriatic lesions. We combined experimental results and network functional analysis to reconstruct the model of *PPARγ* downregulated signaling in psoriasis. We hypothesize that the expression of *IL17, STAT3, FOXP3*, and *RORC* and *FOSL1* genes in psoriatic skin are correlated with the level of *PPARγ* expression and they belong to the same signaling pathway that regulates the development of psoriasis lesion.

## INTRODUCTION

Psoriasis is an example of chronic inflammatory skin disorder with a complex multifactorial origin. Multiple genes cause heterogeneous heredity of psoriasis (Nickoloff and Nestle, 2004; Peters et al., 2000). Interactions of predisposing genes, as well as environmental influences, are required for the development of the disease.

Family genotyping supports the hypothesis that different phenotypes or manifestations of psoriasis are determined by different genetic loci (Samuelsson et al., 1999). These loci are associated with psoriasis and located at least on 13 different chromosomes and are named PSORS (Psoriasis Susceptibility) PSORS1-PSORS13 (Hız Meliha Merve, 2017). Each PSORS contains a list with several revealed genes-candidates (Barker, 2001).

Peroxisome proliferator-activated receptors (PPARs) do not get on lists of gene-candidates for psoriasis, however, the important role of PPARs in anti-inflammatory and immunomodulatory cellular signaling pathways has been established. Recently association of proline12/alanine gene polymorphism (rs1801282) in peroxisome proliferator-activated receptor gamma (*PPARγ*, NCBI Gene ID: 5468) was found to be associated with psoriasis and obesity in Egyptian patients (Seleit et al., 2019).

PPARs perform function primarily as ligand-dependent transcription factors which activate genes with PPAR-responsive elements (PPREs) in their promoter. *PPARγ* is detected mostly in well-differentiated suprabasal keratinocytes within the human epidermis (Icre et al., 2006). Human hair follicle epithelial stem cells also express *PPARγ* which maintains their survival in normal conditions (Billoni, Bruno Buan, Brigitte Gauti, 2000). Skin adipocytes and sebocytes are the next large *PPARγ* depositions (Alestas et al., 2006; Inoue et al., 2014) and the protein is vital for their differentiation (Nehrenheim et al., 2013; Paus et al., 2007). The *PPARγ* expression was reported to be downregulated in the psoriatic skin of mice and DDH1 dose-dependently could restore the gene expression (Kitahata et al., 2018). In vitro experimental models of psoriasis showed the expression of other PPARs (PPARa) was also decreased in the skin, while PPARb and PPARd expression were increased (Friedmann et al., 2005). Mice model of inflammatory skin diseases revealed that the expression of *PPARγ* and PPARa was decreased in the skin due to the absence of the *Dlx3* gene (Hwang et al., 2011). The medical suppression of *PPARγ* improved the health of the mice model of atopic dermatitis (Jung et al., 2011). Wang X at all. reported that gene *PPARγ* had high level of expression in the skin of IMQ-induced psoriasis mice, and a *PPARγ* -selective antagonist GSK3787 was able to decrease the inflammation in the skin (Wang et al., 2016). Finally, another animal model research showed that mutant mice with deleted *PPARγ* did not have sebaceous glands and normal hair follicles (HF), and developed scarring alopecia and skin inflammation (Sardella et al., 2018). There is no experimental evidence about *PPARγ* activity level in human skin of patients with psoriasis to our knowledge.

*PPARγ* signaling in psoriasis has been studied at a good level, but conflicting experimental results do not allow describing a clear picture of protein-protein interactions and pathological changes in cell pathways leading to the development of psoriasis (read below, section “Pathway model of PPARγ signaling in psoriasis”).

In this work we tested a hypothesis that low levels of *PPARγ* may change the activity of cellular signalling pathways in the skin and facilitate the chronic inflammatory and immune response in psoriatic lesion in humans. Based on the literature-based protein-protein interactome (PPI) and pathway analysis we proposed that low *PPARγ* activity promotes the development of psoriatic lesions due to changes in the inflammatory signaling pathways regulated by STAT3, RORC, FOXP3, FOSL1 and IL17A. To check the hypothesis, we measured the expression of these genes altogether with *PPARγ* on the mRNA level in the skin of patients with psoriasis before and after low-intensity laser treatment.

## MATERIALS AND METHODS

### Protein-protein interactome (PPI) analysis and pathway model reconstruction

To reconstruct the *PPARγ*-psoriasis interactome we used the literature-based database PSD (Resnet - 2020 ®, Elsevier Pathway Studio database). PSD is a mammal - centered database where relationships between biological terms and molecules extracted from published papers with natural language processing technology (NLP). Data from public databases with experimental types of connections are also present in PSD. Resnet - 2020 contains over one million objects and more than 12 million relationships with more than 55 million supporting sentences ((Nesterova et al., 2019), www.pathwaystudio.com).

For PPI analysis we used SQL language and ran queries to filter PSD connections and found inhibited by *PPARγ* expression targets that simultaneously have positive relationships with psoriasis (see *“PPARγ* targets and regulators” file, list 1 in supplemental materials). To find *PPARγ* regulators we selected genes that negatively regulate expression of *PPARγ* and simultaneously negatively regulate *PPARγ* expression targets (see *“PPARγ* targets and regulators” file, list 2 in supplemental materials). To focus only on gene expression signaling and exclude other molecular types of interaction, we considered only two types of relationships in PSD that indexed sentences about the changing of mRNA or gene expression (“Expression” and “PromoterBinding). Queries and other parameters of network filtering are available by a request.

We used Pathway Studio software to reconstruct the model of *PPARγ* signaling. Models are interactive networks which describe connections between molecules and related phenotype or biological processes. Models are kept in RNEF format, connected with PSD and include different annotations of molecules and relationships (synonymes, identificators, references, sentences, effects, mechanism of actions and more). All files can be found in supplemental materials (see below).

### Pathway functional analysis

List of proteins that we had identified in the PPI analysis was set up with Sub-Network Enrichment Analysis (SNEA, Pathways Studio), Fisher exact test, Enrichr tool (Chen et al., 2013), and KEGG mapping tool (Kanehisa and Sato, 2020). SNEA was used to find cell processes statistically enriched with genes from list 1 and 2. SNEA is the modification of gene set enrichment analysis that accounts for relationships between genes in the network (Kotelnikova et al., 2010). Fisher test was used to find associated Pathway Studio pathways and Gene Ontology (GO) functional gene groups (Ashburner et al., 2000). Associated KEGG pathways we found with the KEGG mapping tool and other associations we found with Enrichr tool.

Cell processes were selected if more than 5 genes from the list 3 (combined genes from list 1 and list 2) were overlapped with total genes associated with the pathway, and if more than 5% genes from the list 1 and 2 were overlapped with a sub-network or GO group. We selected top sub-networks and KEGG pathways filtered by rank, and top PS pathways and GO groups filtered by Jaccard index. For the comparison of methods, we selected top 50 sub-networks, 50 pathways, and 50 GO groups after manual filtering off unrelated diseases (such as cancer), viral and bacterial KEGG pathways. See supplemental materials for results of pathway functional analysis (“PPARG network analysis” file and “PPARG Enrichr analysis” file).

### Microarray in-silico analysis

Public microarray data (GEO, GSE13355) was used to verify the reconstructed model of *PPARγ* signaling in psoriasis. GSE13355 contains data about the expression of the human genome in skin samples of 58 patients with psoriasis (Ding et al., 2010). DE (differentially expressed genes) were identified with a two-class unpaired T-test between samples of lesional skin of each patient (PP samples) and non-lesional skin uninvolved samples (PN samples). Multiple probes were averaged by the best p-value or maximum magnitude. Pathway Studio software was used for calculation of DE and pathway analysis.

### Supplemental materials

All supplemental materials are available to download from ResearchGate resource by the link https://www.researchgate.net/publication/340427568_Supplemental_Materials_The_role_of_PPARg_downregulated_signaling_in_psoriasis (Sobolev, 2020). All pathways models and their annotations are available for browsing and can be downloaded at http://www.transgene.ru/ppar-pathways.

## RESULTS AND DISCUSSION

### Reconstruction of downregulated *PPARγ* pathway model associated with psoriasis

For testing the hypothesis that low levels of *PPARγ* trigger inflammatory signaling pathways in the skin, we analysed protein-protein interaction literature-based network (PSD, Elsevier Pathway Studio) and several public ontologies and databases (Gene Ontology, Human Protein Atlas, KEGG, Reactome).

First, in the PSD network, we identified *PPARγ* downstream expression targets and upstream regulators (inhibitors) of *PPARγ* expression. For researching the downstream targets, we looked for genes and proteins which were reported to be inhibited by *PPARγ* and simultaneously were positive biomarkers for psoriasis. 146 associated with psoriasis genes and gene families whose expressions are repressed by *PPARγ* had been found. For researching the upstream of *PPARγ* signaling we focused on the transcriptional factors which can inhibit both the expression of *PPARγ* and his direct targets. 99 associated with psoriasis unique negative regulators of *PPARγ* had been identified. Then we combine regulators with targets to obtain 182 names of unique genes forming the *PPARγ* down-regulated sub-network associated with psoriasis (see supplemental materials, “PPARG regulators and targets” file, list 3). (Figure 1).

**Figure 1.**
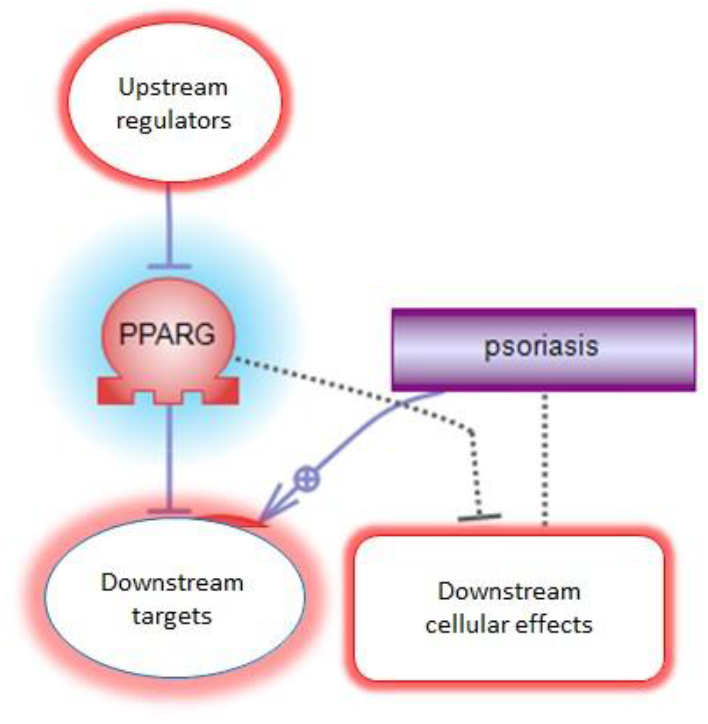
The logic of discovering the members of *PPARγ* down-regulated sub-network associated with psoriasis.

### Comparative pathway analysis of PPARγ downregulated signaling associated with psoriasis

Several methods of pathway analysis were performed to explore the functional roles of 182 targets and regulators of *PPARγ* revealed in PPI analysis. Methods of pathway functional analysis are widely used for discovering cellular processes and signalings that are statistically associated with the list of genes or proteins (Nesterova et al., 2020).

We compared results from pathway functional analysis with three resources: Gene Ontology, Elsevier Pathways, and KEGG Pathways. Gene Ontology is the source of groups of proteins or genes manually assigned by their different functional roles. Elsevier Pathways and KEGG Pathways are manually reconstructed schemas or models of interactions between proteins describing molecular mechanisms of one or several biological processes. Gene Set Enrichment Analysis (GSEA) is a well-known method to analyse predefined and manually created collections of gene groups and pathways (Subramanian et al., 2005). Besides GSEA we used SNEA method which allows finding associated cellular processes based on literature - based PPI network. SNEA does not use predefined groups of genes or pathways and is considered less biased (Kotelnikova et al., 2010; Nesterova et al., 2020).

According to the results of comparative pathway analysis, *PPARγ* downregulated signaling is associated with adipogenesis, activation of myeloid pro-inflammatory cells (with a predominance of mast cells and dendritic cells), and activation of overall immune system response (with a predominance of Th17 cells). Also, fibrogenesis, cell-to-cell contacts, vascular-related processes and universal cell processes, such as cell proliferation or cell death, were identified (Figure 2). Cellular possesses directly associated with psoriasis were present in results from each source that we compared (Table 1).

**Figure 2.**
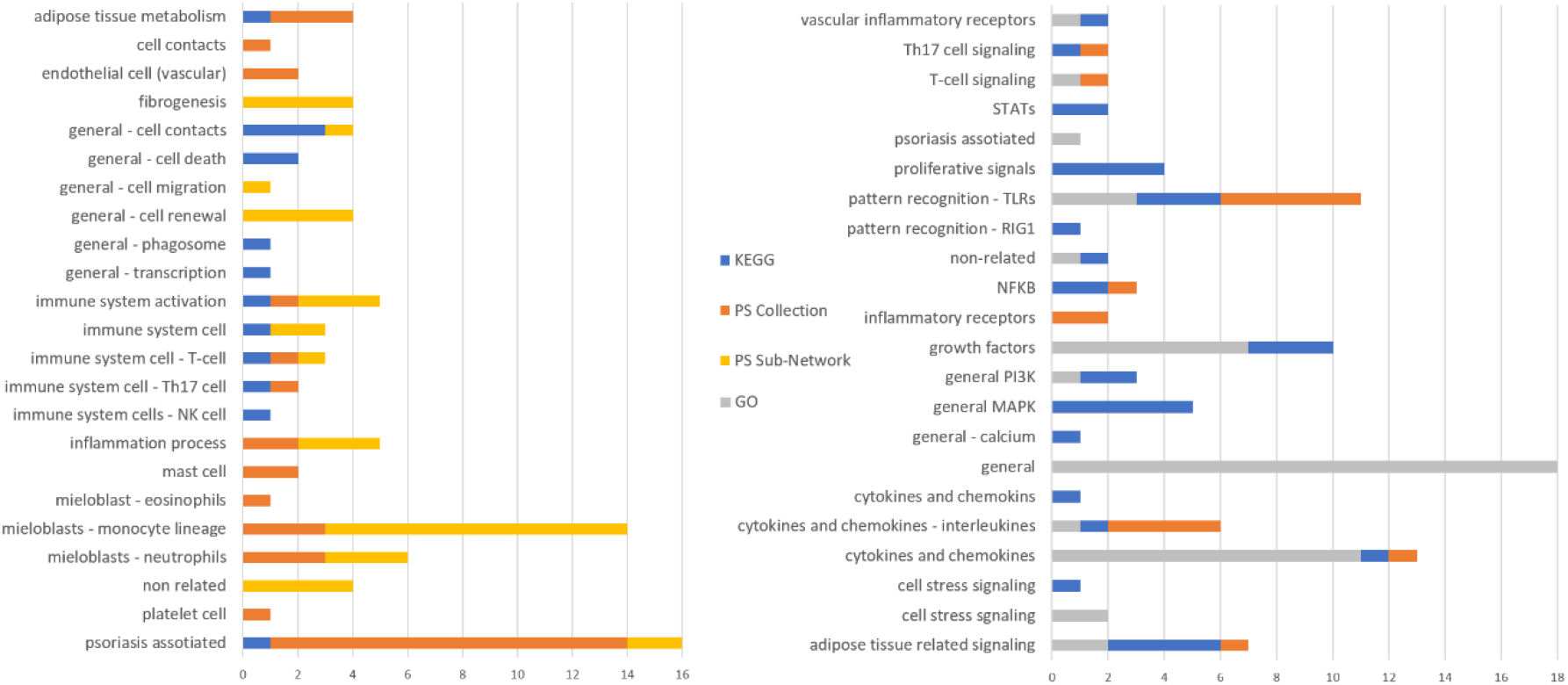
Cellular processes associated with members of PPARγ down-regulated signaling associated with psoriasis. Numbers are for the sum of pathways, subnetworks or GO groups in each category. Different sources are highlighted with blue (KEGG database), orange (Pathway Studio Pathway Collection), light orange (Resnet-2020 database), and grey (GO). For the complete list of results, see supplemental materials, “PPARG network analysis” file).

**Table 1.** Cell processes directly related to psoriasis and enriched with genes and proteins from the *PPARγ* downregulated signaling. See complete results with additional statistics in supplemental materials, “PPRAG network analysis” file.

| Name of the process or signaling | Source | Rank or Jaccard similarity (the closer to 10%, the more similarity) | Category |
| --- | --- | --- | --- |
| Keratinocyte Activation in Psoriatic Arthritis | PS Collection | 9.13% | Disease |
| lesion size | PS Sub-Network | 46 | Targets neighbors |
| keratinocyte proliferation | PS Sub-Network | 100 | Targets neighbors |
| Skin Fibrosis | PS Collection | 9.09% | Disease |
| T-Cells Differentiation Block in Psoriasis | PS Collection | 8.88% | Disease |
| Th17-Cell and Th1 Immune Response in Psoriatic Arthritis | PS Collection | 8.82% | Disease |
| Dendritic Cell Dysfunction in | PS Collection | 8.44% | Disease |
| Psoriatic Arthritis |  |  |  |
| T-Cell Cytotoxic Response against Melanocytes in Vitiligo | PS Collection | 6.20% | Disease |
| Synovial Fibroblast Activation in Psoriatic Arthritis | PS Collection | 8.18% | Disease |
| Inflammatory Reaction in Acne Vulgaris | PS Collection | 5.70% | Disease |
| Apoptotic Keratinocytes Clearance Recession in Systemic Lupus Erythematosus | PS Collection | 5.69% | Disease |
| Atopic Dermatitis | PS Collection | 5.62% | Disease |
| Hair Follicle Keratinocyte Apoptosis | PS Collection | 5.56% | Disease |
| Vitiligo | PS Collection | 5.07% | Disease |
| Melanogenesis - Homo sapiens | KEGG | n/a | Pathway |
| positive regulation of timing of anagen | GO | 1.10% | GO: biological_process |
| glycosaminoglycan binding | GO | 4.24% | GO: molecular_function |

Top sub-networks from SNEA were neighbours of adipogenesis and adipocyte differentiation, followed by the immune response, and T-development. The sub-networks “neighbours of monocyte recruitment or differentiation” and “macrophage differentiation” had the most percent (9%) of overlapped genes from *PPARγ* down-regulated signaling.

GSEA analysis of PS Pathway Collection and KEGG pathways resulted in many cancer-related processes. The disease taxonomy filtering with PS pathways about skin and immune system identified processes related to adipokines and IL17 signaling (Table 2). The signaling of aryl hydrocarbon receptor (AHR) in Th17 cells was the pathway with the biggest percent (48%) of overlapped genes from *PPARγ* down-regulated signaling.

**Table 2.** List of PS pathways associated with members of PPARγ down-regulated signaling associated with psoriasis. See complete results with additional statistics in supplemental materials, “PPRAG network analysis” file.

| Pathway name | # of Entities | Overlap | Percent Overlap | p-value | Jaccard similarity | Pathway Taxonomy Top |
| --- | --- | --- | --- | --- | --- | --- |

|  |  |  |  |  |  | Category |
| --- | --- | --- | --- | --- | --- | --- |
| EGFR -> Expression Targets in Skin | 98 | 26 | 26 | 4.68E-11 | 10.28% | Biomarkers |
| GPCRs Family -> Expression Targets in Lymphoid System and Blood | 89 | 24 | 26 | 1.97E-10 | 9.76% | Biomarkers |
| Adipokines Production by Adipocyte | 58 | 20 | 34 | 4.8E-16 | 9.13% | Biological Process |
| Skin Fibrosis | 83 | 22 | 26 | 3.29E-17 | 9.09% | Disease |
| Scavenger Receptor OLR1 in Inflammation Related Endothelial Dysfunction | 73 | 21 | 28 | 2.8E-17 | 9.01% | Disease |
| T-Cells Differentiation Block in Psoriasis | 52 | 19 | 36 | 5.87E-18 | 8.88% | Disease |
| Adipokines Production by Adipocyte Impaired in Obesity | 56 | 19 | 33 | 2.99E-17 | 8.72% | Disease |
| CD40 -> Expression Targets in Thymus | 58 | 19 | 32 | 4.67E-10 | 8.64% | Biomarkers |
| Lymphocyte Mediated Myocardial Injury in Myocarditis | 84 | 21 | 25 | 6.67E-16 | 8.61% | Disease |
| IL17 Signaling in Psoriasis | 49 | 18 | 36 | 4.08E-17 | 8.49% | Disease |
| Th17-Cell Differentiation | 73 | 19 | 26 | 8.94E-13 | 8.09% | Biological Process |

Among top KEGG pathways enriched with our gene list, we identified general MAPK and PI3K signaling and cancer-related pathways (for example, “hsa05200 Pathways in cancer - Homo sapiens”). TNF signaling pathway (hsa04668), as well as Th17-cell (hsa04659) and IL17 pathway (hsa04657), were also in the top 10 results. The cytokine-cytokine receptor interaction (hsa05200) and PI3K-Akt signaling (hsa04151) had the highest number of overlapped entities (48 and 34).

The list of revealed in pathway analysis molecular cascades complete the lists of cell processes.

There was no surprise that activation of general cellular flows like ERK/MAPK, RAS/ACT1, and adipose cells related AMPK, mTOR, and cAMP cascades were associated with the list of *PPARγ* targets and regulators. Also, among the top of associated molecular signalings there were well predictable inflammatory cascades like Toll-like receptors, interleukins and interleukins receptors signaling (IL17, IL1B, IL6, and IL1R1) altogether with all-purpose cytokines and cytokines receptors signaling (CXCR3, CCR1, TNF). Signalings related to transcription factors NFKB and STATs also were significantly associated with the analysed list. GO functional group “GO: glycosaminoglycan binding”; “IL1R1 signaling in Pneumocytes’’ from PS Pathway Collection; and “ErbB signaling pathway” (hsa04012) from KEGG had the maximum rank (See complete results with additional statistics in supplemental materials, “PPRAG network analysis” file). Glycosaminoglycans are essential for skin functioning. IL1R1 is a receptor commonly activated in any non-specific inflammatory processes. Finally, the ERbB/EGFR family is involved in cell proliferation and tumor development.

Additional comparison of pathway analysis results with other pathways databases (WikiPathways, Reactome, Biocarta analysed with Enrichr tool) confirmed results obtained with PS Pathway Collection (Figure 3). Pathways from all sources revealed skin inflammatory processes, TLRs and interleukins related cascades. However, the list of molecules was different compared with PS and KEGG results presenting IL10 and IL22R and no IL17 associations. In addition, analysis with DisGeNET (Piñero et al., 2017) confirmed that the *PPARγ* regulators and targets are connected with psoriasis since top diseases associated with the list 3 were: psoriasis, epithelial hyperplasia of skin, and inflammatory dermatosis. Allergic reaction, neutrophilia, and vascular diseases were also in the top 10 results (see supplemental materials, file “PPARG Enrichr analysis” file).

**Figure 3.**
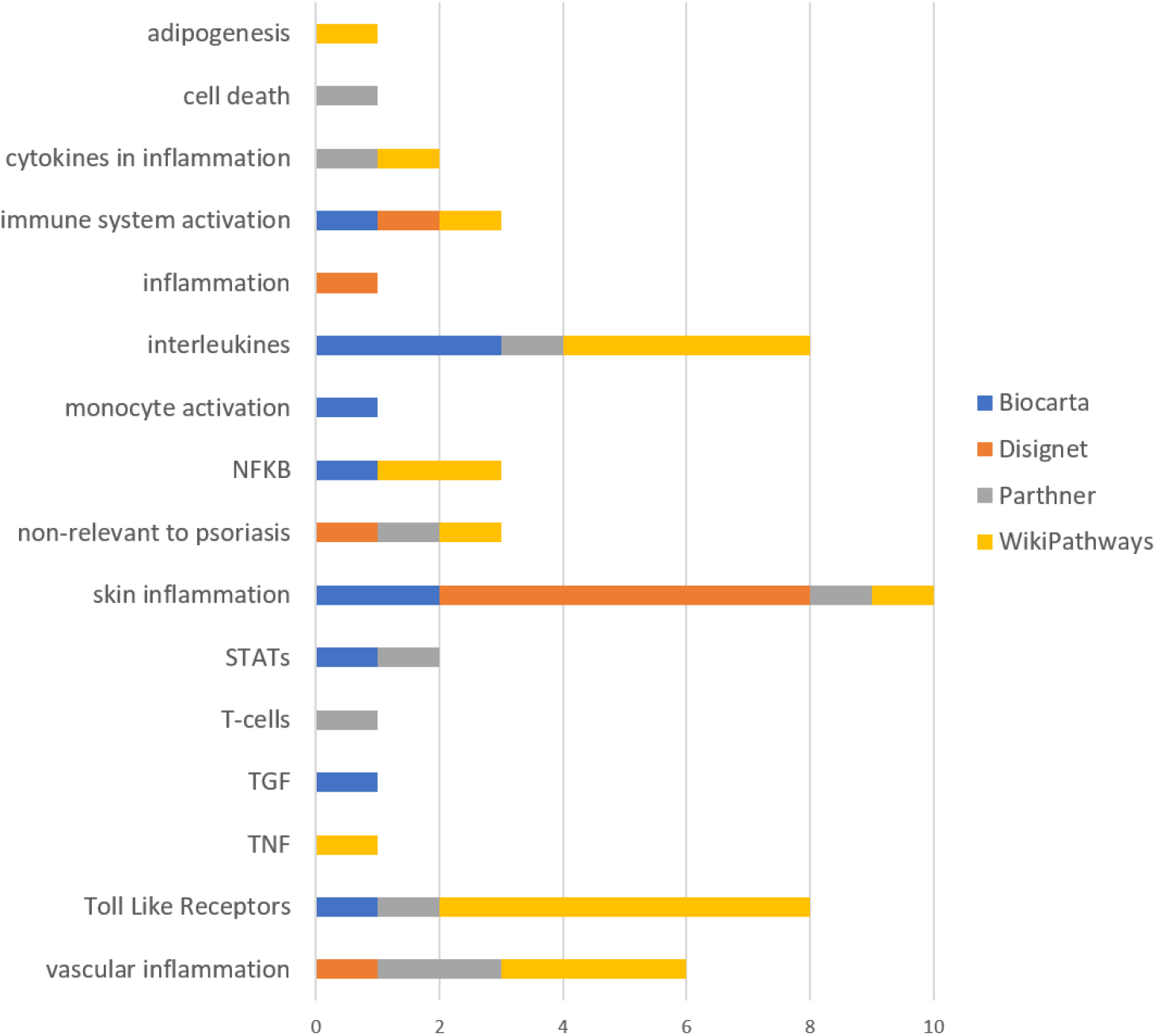
Comparison of results of pathway analysis (GSEA) with different sources for PPARγ down-regulated signaling associated with psoriasis. Results were calculated with the Enrichr tool. Numbers are for the sum of pathways in each category. Different sources are highlighted with blue (Biocarta database), grey (Partner database), orange (DisGeNET) and yellow (WikiPathways).

### Pathway model of PPARγ signaling in psoriasis

Considering results of PPI network and functional pathway analysis we build a hypothetical model that describes cellular molecular mechanisms of involvement of *PPARγ* in the maintenance of chronic inflammatory and immune response in human psoriatic skin. Literature - based network (PSD) were used to build the model. Figure 4 described the adopted for the publication simplified version of the downregulated PPARγ pathway model. See supplemental materials for the completed version of the pathway model.

**Figure 4.**
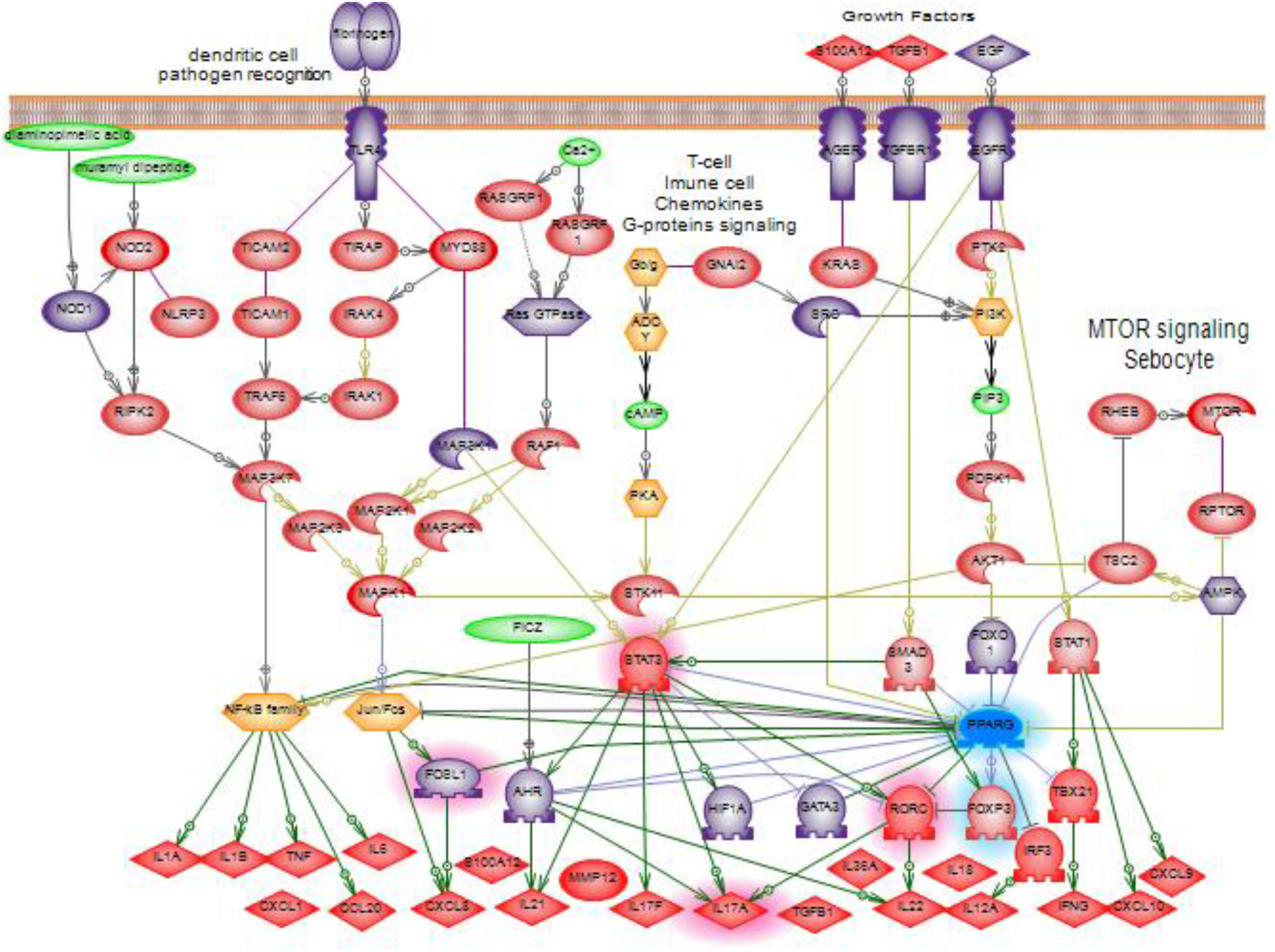
Model of downregulated *PPARγ* signaling in psoriasis. *PPARγ* which expression is downregulated in psoriasis is colored blue. Regulators that may inhibit the expression of *PPARγ* (data based on PPI and pathway functional analysis) are colored in violet. Targets which may be over-expressed in psoriasis due to the decrease of the negative impact of *PPARγ* are colored in bright red. *FOXP3, STAT3, ĨL17A, RORC* and *FOSL1* are highlighted according to own experiments (read section “Preliminary analysis of *PPARγ* signaling in human psoriatic skin”). Down-expressed genes (*PPARγ, FOXP3*) are highlighted in blue, overexpressed are highlighted in red.

Based on the model, reducing the level of the *PPARγ* gene expression may be a result of the over-regulation of several cascades. Pattern recognition receptors (TLRs, NOD1, NOD2, CLEC7A) that sensor pathogens and highly expressed in keratinocytes and monocytes during the infection may be one of such cascades. All-purpose cellular cascades like growth factors signaling, G-proteins and MTOR signaling also were reported to be inhibitors of *PPARγ* expression in literature and revealed in our analysis. Moreover, transcription factors including NF-kBs, JUN-FOS, AHR, GATA3, HIF1A, FOXO1 and FOSL1 can directly inhibit *PPARγ* expression. All these transcription factors are over-stimulated in the inflammatory and immune response. For example, it is reported that NF-kBs are stimulated in systemic inflammatory processes in general, and in psoriasis as well (Tang et al., 2010; Xu et al., 2015).

In healthy skin *PPARγ* inhibits mentioned transcriptional factors in a feedback regulation loop. *PPARγ* may directly bind and suppress transcriptional factors STAT3 and RORC, by thus blocking the synthesis of pro-inflammatory cytokines including IL17. Less quantity of expressed cytokines decreases the Th17 cell proliferation, minimises chemotaxis of neutrophils and monocytes and results in the reduction of inflammation in psoriatic lesions.

IL17 which is produced mostly by TH17 cells plays the central role in the development of psoriasis because it stimulates keratinocytes to secrete pro-inflammatory cytokines and anti-bacterial peptides (Srivastava et al., 2017). IL17 pathway and Th17 cells had a strong association with *PPARγ* downregulated signaling confirmed by our network and functional analysis.

Th17 cells need robust activity of STAT3 gene for their function and differentiation. Also, STAT3 is described as an important linkage between keratinocytes and immune cells (Chowdhari and Saini, 2014). Previously the expression of *STAT3* was shown to be repressed due to *PPARγ* activation (Hsu et al., 2016). *STAT3* may also act as a regulator of *PPARγ* expression however it is not clear whether with positive or negative effect (Tuna et al., 2014).

As a transcription factor, STAT3 is reported to be a strong activator of RORC (ROR*γ*) and, probably, IL17 gene expression. From the other side, gene RORC is the major inductor of the expression of IL17 cytokines family (Takaishi et al., 2017). *PPARγ* was shown to bind the *RORC* promoter and suppress its expression altogether with *RORC*-mediated Th17 cell differentiation (Hermann-Kleiter et al., 2012).

Transcription factor FOXP3 is closely associated with psoriasis and the diminishing of Treg-cell number (Jorn Bovenschen et al., 2011; Shu et al., 2017). It was shown that activated *PPARγ* induces the stable *FOXP3* expression by strong inhibiting effect on DNA-methyltransferases. The activating effect of *PPARγ* on FOXP3 results in the proliferation of iTreg-cells (Lei et al., 2010).

*FOSL1* is the transcriptional factor which plays important role in many processes related to cell differentiation and tissue remodeling (Sobolev et al., 2011, 2010; Young and Colburn, 2006). *FOSL1* (FOS-like antigen 1) is expressed in low level in healthy tissues, however its expression rises due to presence of mitogens or toxins. The accumulation of the FOSL1 protein in the skin depends on the stage of the keratinocyte differentiation (Mehic et al., 2005). Markers of stratum corneum differentiation like gene IVL are the main expression targets of FOSL1 (Adhikary et al., 2004).

The degree of the pathogenicity of downregulated *PPARγ* in psoriatic lesion depends on the cell type. It is known that *PPARγ* is expressed in Th17 cells as well as in keratinocytes, sebocytes and other cells of the psoriatic lesion (Billoni, Bruno Buan, Brigitte Gauti, 2000; Icre et al., 2006; Inoue et al., 2014; Nehrenheim et al., 2013; Paus et al., 2007). Functional and network analysis supported the association of *PPARγ* down-regulated signaling with keratinocytes, vascular endothelium, vascular smooth muscle cells, macrophages, fibroblasts and adipocytes, and monocytes lineage (particular with CD33+, CD14+ monocytes) (Figure 5). However, we did not attempt to separate the *PPARγ* pathway model by appropriate cell types which is a disadvantage of this work. There is no reliable way to take in account cell specificity in our modeling paradigm. Moreover, we expect that most of the revealed from the literature network analysis cascades will be equal for different human cells due to insufficient experimental studies.

**Figure 5.**
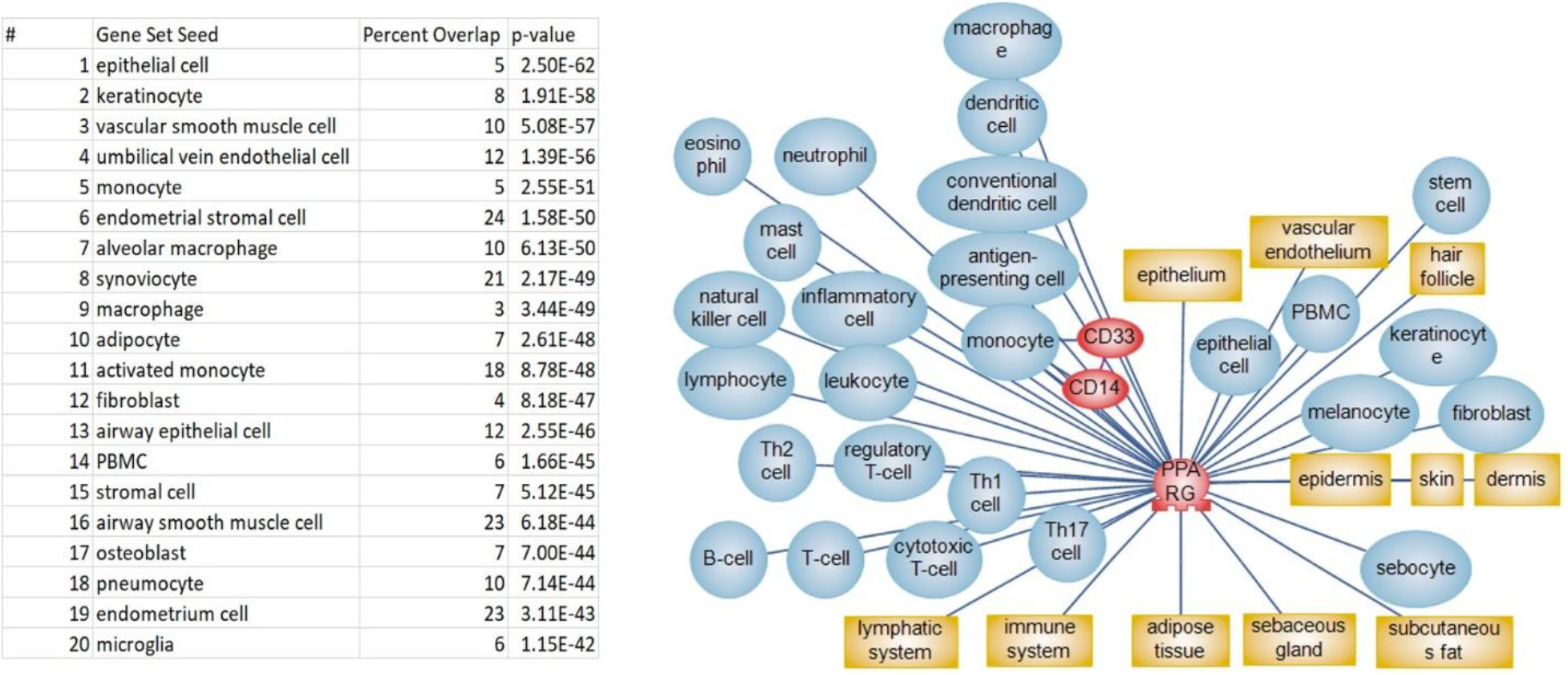
Cells associated with downregulated *PPARγ* signaling. SNEA method and Pathway Studio were used to calculate the results. See the complete list with statistics in supplemental materials, “PPARG and cell” file.

For additional evaluation of the reconstructed model, we analysed the public microarray data (GEO:GSE13355). In that experiment biopsies from 58 psoriatic patients were run on Affymetrix microarrays containing more 50 000 gene probes (Nair et al., 2009). We uploaded raw data from GEO and calculated differentially expressed genes (DE) between samples of psoriatic skin and unaltered samples for all patients. Then we used pathway analysis to explore the difference in the expression for genes of the *PPARγ* model we build (Figure 6).

*PPARγ* gene was slightly down regulated in psoriatic lesions comparing to non-altered lesions in GSE13355 microarray data (Figure 6, *PPARγ* expression diagram).

**Figure 6.**
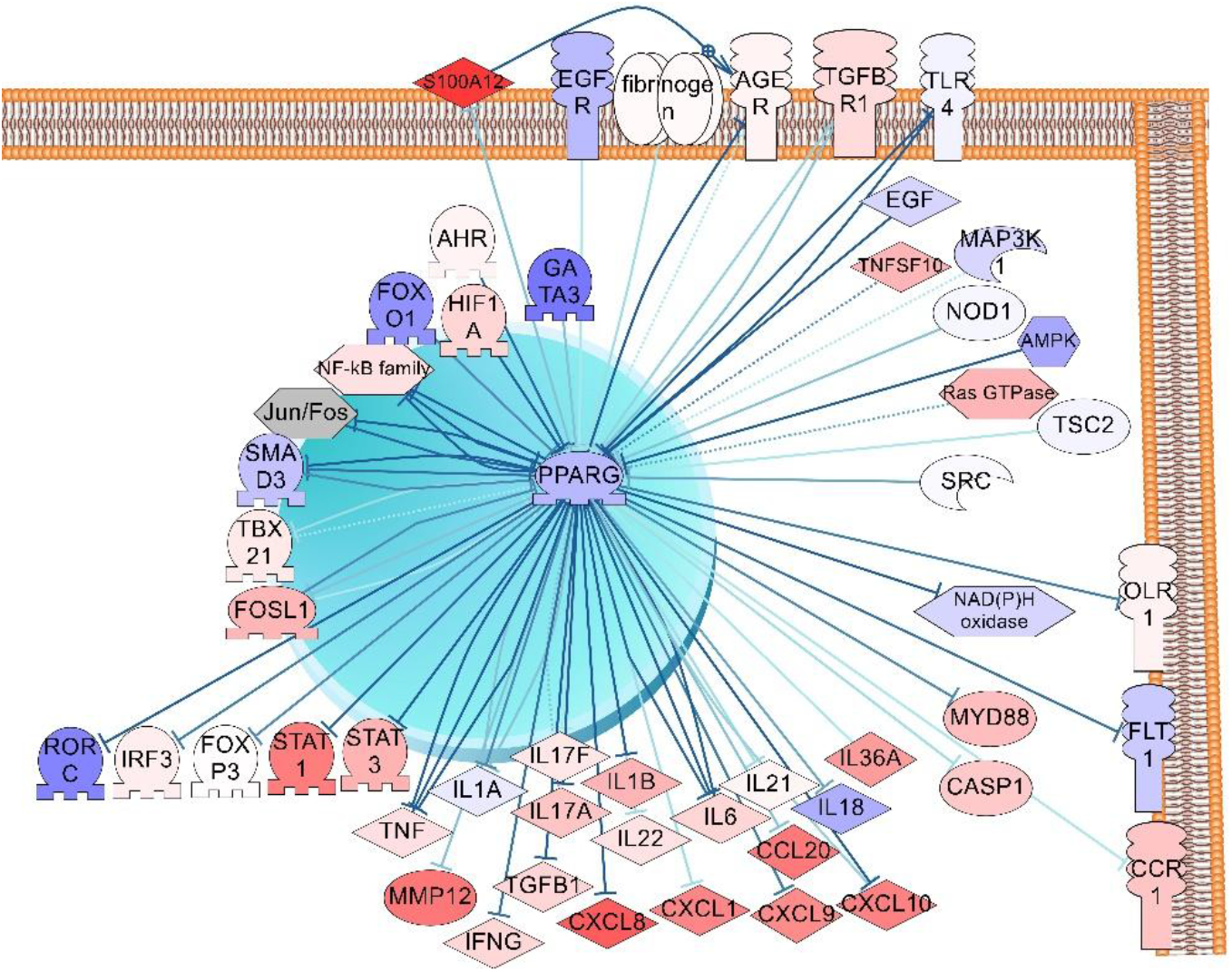
Evaluation of *PPARγ* downregulated sub-network (selected *PPARγ* regulators and targets) associated with psoriasis using microarray data analysis (results of differential expression analysis of psoriatic lesions vs unaltered lesions). The saturation in blue indicates the degree of genes down-expression in psoriatic samples in comparison with unaltered samples. The saturation in red indicates the degree of genes over-expression. The list of targets and regulators see in supplemental materials, “PPARG regulators and targets”, list 3.

We assumed that regulators of *PPARγ* signaling should have higher expression in psoriatic lesion than in normal skin. Only S100A12 (S100 calcium binding protein A12) had a significantly higher level of expression in analyzed microarray data comparing with all regulators of *PPARγ* that we selected for the model (Figure 6, 7). S100A12 binds to the AGER receptor which belongs to the immunoglobulin superfamily and is involved in many processes of inflammation and immune response. S100A12 is thought to be the most prominent biomarker of psoriasis (Wilsmann-Theis et al., 2016). Also, polymorphisms in AGER receptor were found to be associated with psoriasis (Puig and López-Ferrer, 2017).

**Figure 7.**
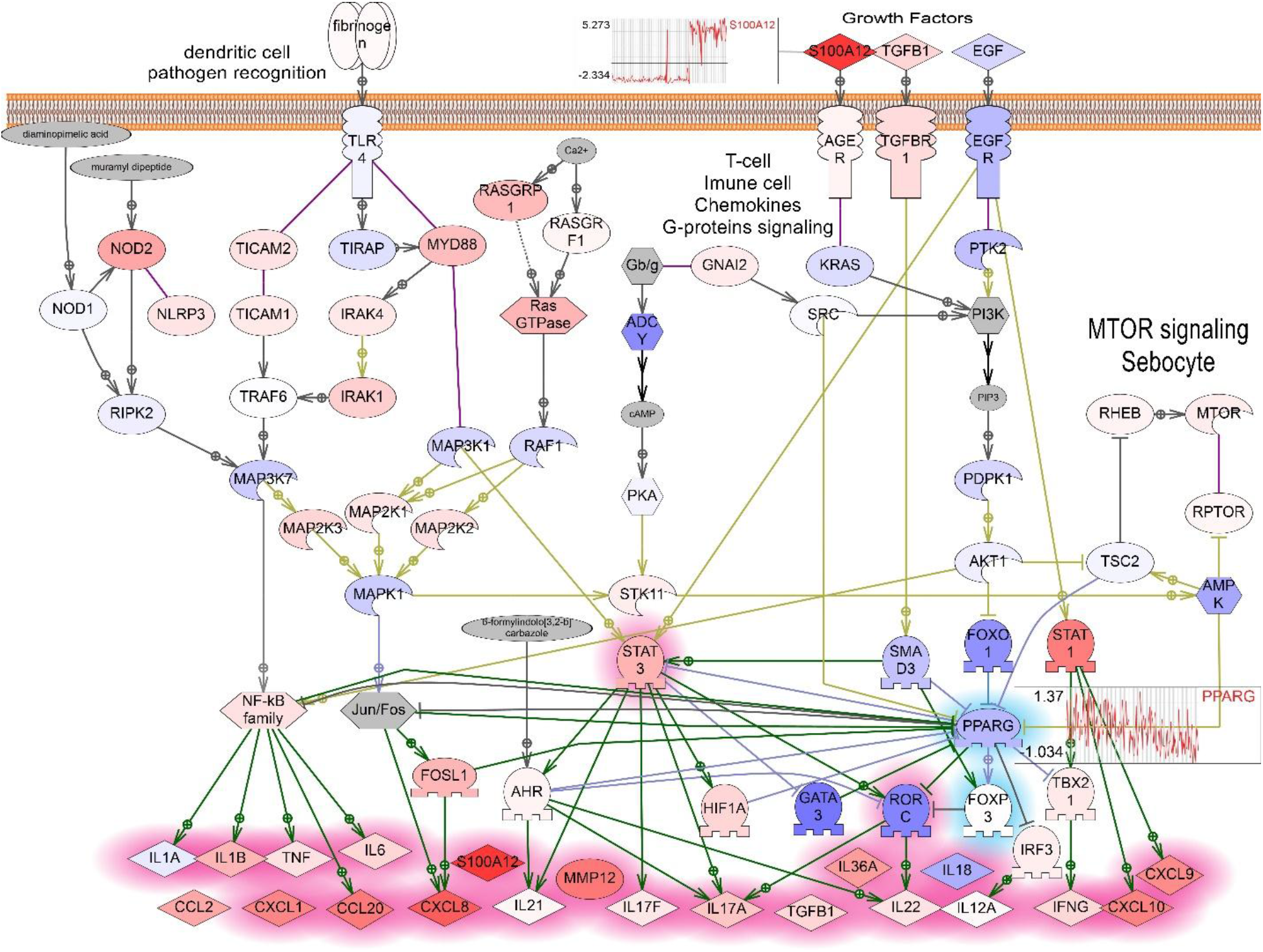
Evaluation of *PPARγ* downregulated model associated with psoriasis using microarray data analysis (results of differential expression analysis of psoriatic lesions vs unaltered lesions from GEO:GSE13355). The saturation in blue indicates the degree of genes down-expression in psoriatic samples in comparison with unaltered samples. The saturation in red indicates the degree of genes over-expression. The plots of expression pattern in psoriatic lesions compared with healthy skins samples are shown for *PPARγ* and *S100A12* genes.

The EGFR signaling almost completely was down-expressed in this microarray data including the *FOXO1* expression which is one of the direct inhibitors of *PPARγ*. Therefore, EGFR / FOXO1 signaling probably does not play an important role in the regulation of *PPARγ* in psoriasis (Figure 7).

## CONCLUSION

In our previous work, we reviewed the recent progress in psoriasis pathways and published two pathway models. The first pathway model described the shift to TH17 cell production during the differentiation of psoriatic T cells. The primary cause of the shift of the T-cell differentiation is supposed to be genetic mutations, for example in IL23R receptors. The second model showed how elevated levels of IL17 and IL22 may activate keratinocytes to release different cytokines and chemokines for attracting neutrophils and other inflammatory cells in the psoriatic lesion (Nesterova et al., 2019).

In this work, we tested the hypothesis that *PPARγ* signaling when downregulated may promote psoriasis. We built the model of *PPARγ* dependent pathways involved in the development of the psoriatic lesions. However, we used a different approach for reconstructing the pathway model and selected key members with bioinformatic analysis. We included in the pathway model top statistically significant regulators of *PPARγ* gene expression and *PPARγ*-depended expression targets. Then we included significant molecular cascades and cell processes from results of the functional analysis (IL17 signaling, TLRs signaling, activation of STAT3 or NFKB transcription factors and others). We tested the model with analysis of published microarray data.

While the prominent role of *RORC* in psoriasis as the major controller of Th17 cell differentiation is well described, however, the evidence of *RORC* expression in psoriasis is controversial and supported by work where mice T-cells and dendritic cells had increased STAT3/RORC expression (Nadeem et al., 2017) still patients with psoriasis had elevated level of *RORC* (RORG-t isoform) (Mendoza et al., 1989). In analysed published microarray data, the level of expression of *RORC* was downregulated in most of 58 patients.

We detected down-regulation of *PPARγ* gene expression in human psoriatic skin from 23 patients with real-time PCR method (data not shown). Our results are similar to data from microarray on 58 patients where average *PPARγ* gene expression also is slightly downregulated in psoriatic lesions (Nair et al., 2009). Our results do not confirm the work of Westergaard M et al. which described the slightly higher level of the *PPARγ* expression in human psoriatic skin compared to normal skin. However, the level of *PPARγ* mRNA was close to the detection limit in their research (Lei et al., 2010). This difference may be due detection of different isoforms of *PPARγ* which all have different patterns of the expression (Meirhaeghe and Amouyel, 2004). More research on protein level is needed to conclude whether *PPARγ* gene expression is downregulated in psoriatic lesions.

Within the framework of the model validation we hypothesize that signalling related to repressed *PPARγ* activity is correlated with the development of psoriasis. IL17A, STATS3, and RORC (RORg) are statistically significant *PPARγ* negative targets and we detected higher levels of their mRNA in psoriatic lesion of 23 patient, and moreover, the decrease of their expression levels after laser treatment (preliminary results, not shown). The alignment of our preliminary experimental results with microarray data and PPI network analysis shows that the reconstructed model of *PPARγ* downregulated signaling in psoriasis can be useful for further research.

## ACKNOWLEDGEMENTS AND FUNDING

We thank the “University Diagnostic Laboratory” LLC for funding along with Elsevier Inc. and Pathway Studio® for software and the database.

